# Super-Resolution NMR Spectroscopy using the Intersection of Non-Redundant Information on Resonance Groups

**DOI:** 10.1101/2022.04.11.487656

**Authors:** Justin L Lorieau, Alexander Malooley, Indrani Banerjee

## Abstract

The resolution of spectra is a major limitation in the application of Nuclear Magnetic Resonance (NMR) spectroscopy to large and complex molecular systems. In this report, we introduce a technique to enhance the resolution of NMR spectra beyond the intrinsic limitations of a spectrometer for a single spectrum by using the Intersection of Non-Redundant Information on Resonance Groups (INIR). With INIR, we reconstruct 900-MHz (21.1T) spectra from a 500-MHz (11.7T) NMR spectrometer, which compare favorably to experimental 900-MHz spectra. INIR holds promise in significantly enhancing the resolution of NMR spectra and in extending the size and complexity of molecules studied by NMR.

## Introduction

A major endeavor in NMR spectroscopy is the increase of magnetic field strengths. Stronger magnetic fields are desirable because they provide greater sensitivity and increased resolution in spectra. Without further advances in semi-conducting materials, it is becoming increasingly difficult and, in some cases, prohibitively expensive to manufacture spectrometers with stronger magnets.

Many alternative approaches exist to increase the sensitivity of NMR experiments: hyperpolarization and cooled samples,^1^ cooled electronics^2^ and improved pulse sequences.^3–5^ Likewise, increased resolution is desirable because it decreases spectral ambiguities and increases the molecular size and complexity of systems studied by NMR. Higher magnetic fields improve resolution by increasing the dispersion of peaks, from the field-dependent chemical shift, while only modestly increasing linewidths for nuclei that have small relaxation contributions from the chemical shift anisotropy (CSA). Alternative approaches for resolution enhancements exist with isotope labeling strategies,^6^ sample preparation methods and pulse sequences.^7,8^ Resolution nevertheless remains an important bottleneck in the development and application of NMR spectroscopy to complex molecules such as proteins and nucleic acids.

In this report, we develop the Intersection of Non-redundant Information on Resonance groups (INIR) to significantly enhance the resolution of NMR spectra. INIR utilizes highly correlated yet non-redundant spectra to accurately define the properties of spins such as chemical shifts, J-couplings, linewidths and dynamic parameters. Conceptually, the range of these parameters can be represented by a circle from a Venn diagram. The incorporation of multiple non-redundant spectra produces a series of partially overlapping Venn diagram circles in which the intersection more accurately defines the parameters of a system.

In this first demonstration, we used an interleaved ^15^N heteronuclear combined quantum coherence (^15^N-HCQC) spectrum to construct 3 highly correlated spectra for the single-quantum (SQ), zero-quantum (ZQ) and double-quantum (DQ) transitions of ^1^H-^15^N resonance groups. A diagram for spectra of these 3 experiments is shown in **Figure 1**. The cluster of 3 peaks arrange in different patterns between SQ, ZQ and DQ transitions, and the linewidths are different between the spectra. Importantly, the features of each cluster of peaks are different between the 3 spectra. In this example, the blue peak’s position and lineshape are more easily resolved in the ZQ spectrum whereas the red peak is more easily resolved in the DQ spectrum. By simulating the resonance groups for all 3 spectra simultaneously, we can accurately define their peak positions and linewidths. Thereafter, we can reconstruct a standard ^15^N heteronuclear single-quantum coherence (HSQC) 2D spectrum with enhanced resolution.

**Figure 1.**
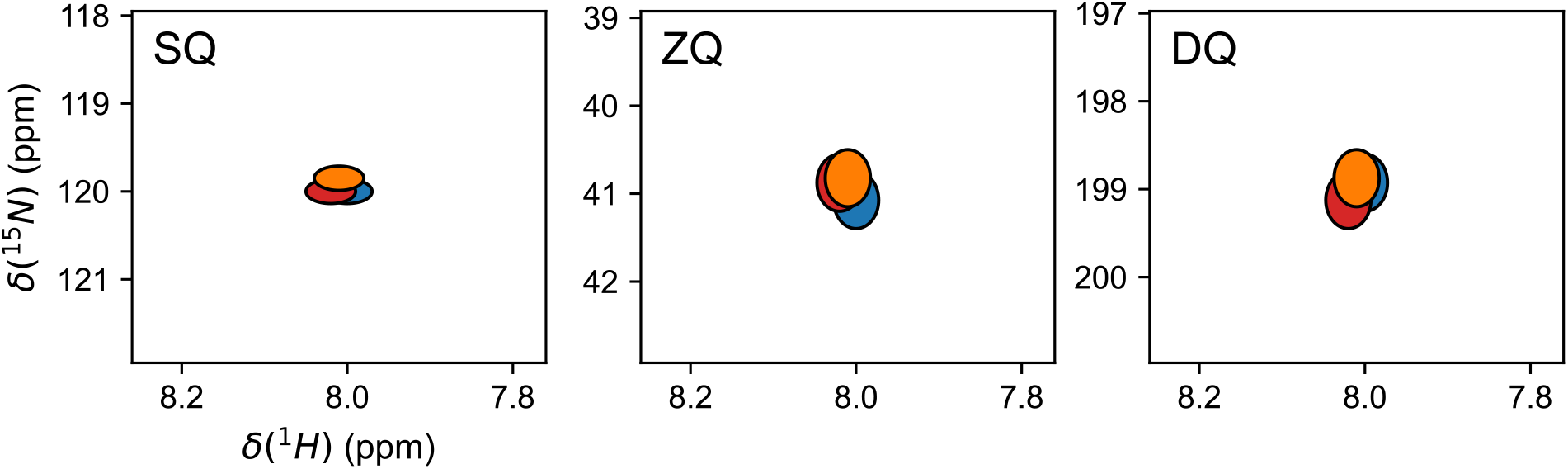
Simulated spectra with a cluster of 3 peaks (ellipses) in blue, red and orange. The SQ spectrum simulates a 500-MHz HSQC with a blue peak (8.00 ppm, 120.00 ppm), a red peak (8.02 ppm, 120.00 ppm) and an orange peak (8.01 ppm, 119.85 ppm). The SQ peaks have a width of 0.06 ppm in the ^1^H dimension and 0.03 ppm in the ^15^N dimension. The ZQ spectrum simulates a 500-MHz HZQC with a blue peak (8.00 ppm, 41.08 ppm), a red peak (8.02 ppm, 40.88 ppm) and an orange peak (8.01 ppm, 40.82 ppm). The ZQ peaks have a width of 0.055 ppm in the ^1^H dimension and 0.66 ppm in the ^15^N dimension. The DQ spectrum simulates a 500-MHz HDQC with a blue peak (8.00 ppm, 198.93 ppm), a red peak (8.02 ppm, 199.12 ppm) and an orange peak (8.01 ppm, 198.88 ppm). The DQ peaks have a width of 0.055 ppm in the ^1^H dimension and 0.66 ppm in the ^15^N dimension. The peak linewidths were modeled against the average full-widths and half-heights of HSQC/HZQC/HDQC peaks from a α-synuclein HCQC spectrum.

With an HCQC experiment collected on a 500-MHz NMR spectrometer (11.7 T), we have simulated and enhanced the resolution of ^15^N-HSQC 2D spectra for two protein systems: α-synuclein and the C-domain of the influenza M1 protein (M1C). We combined the information of 3 spectra to reconstruct an enhanced 500→900-MHz HSQC 2D spectrum for comparison with an HSQC 2D spectrum collected at 900-MHz (21.1 T). We found that the number of assignable peaks was significantly greater for the 500→900-MHz HSQC in comparison to a 500-MHz HSQC, and that this number was comparable to the corresponding 900-MHz HSQC spectrum. The implementation, advantages, drawbacks, and challenges of INIR NMR spectroscopy are discussed.

## Theory

*Pulse sequence*. The HCQC pulse sequence collects interleaved SQ and ZQ/DQ spectra. The first interleaved experiment is a heteronuclear single-quantum coherence (HSQC) with sensitivity improvement and the second interleaved experiment is a ZQ/DQ-resolved heteronuclear multiple-quantum coherence (HMQC) spectrum with sensitivity improvement ^4,9^. The objective of this pulse sequence is to collect the two spectra as closely as possible in terms of pulse sequence elements and experimental time. The resulting HCQC spectrum contains SQ (2H_z_N_+/-_), ZQ (2H_+_N_-_, 2H_-_N_+_) and DQ (2H_+_N_+_, 2H_-_N_-_) coherences (**Figure 2**).

**Figure 2.**
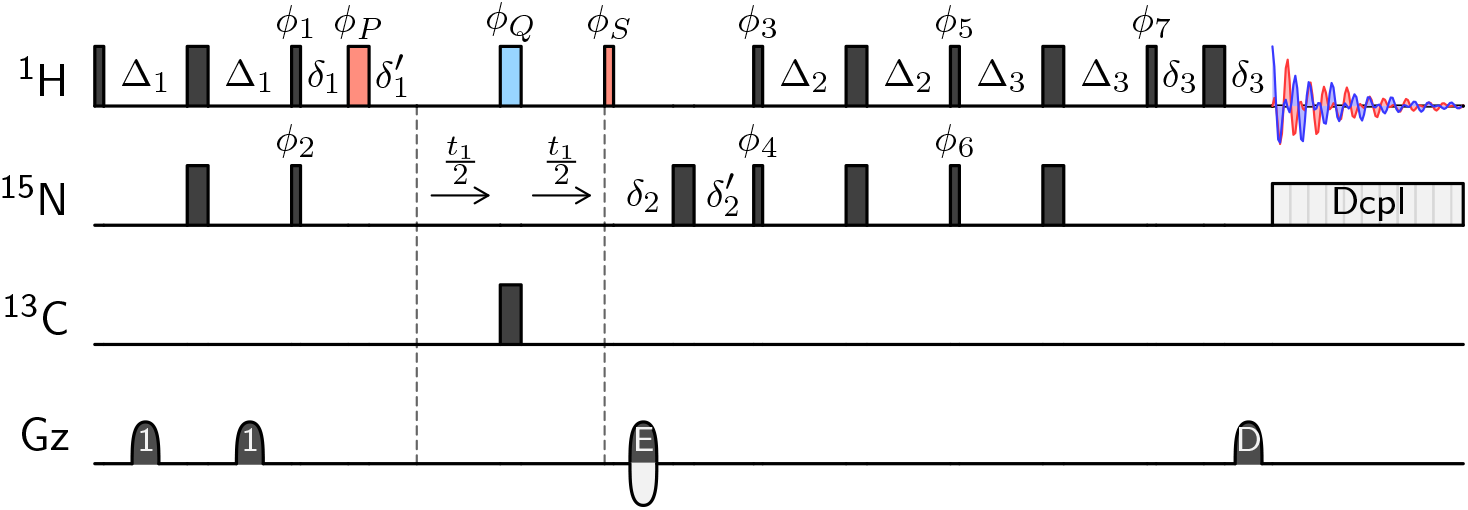
The heteronuclear combination quantum coherence (HCQC) pulse sequence with sensitivity improvement. The pulse sequence is run in interleaved mode for single-quantum (SQ) and zero-/double-quantum (ZQ/DQ) experiments. The thin rectangles represent 90-degree pulses and the wide rectangles represent 180-degree pulses. For the SQ experiment, the blue pulse is active and the red pulses and δ_1_^’^/δ_1_ periods are inactive. The SQ phases are *ϕ*_1_ = y, *ϕ*_3_ = x, *ϕ*_5_ = y, *ϕ*_7_ = x and *ϕ*_Q_ = x. For the ZQ/DQ experiment, the red pulses and δ_1_/δ_1_’ periods are active and the blue pulse is inactive. The ZQ/DQ phases are *ϕ*_1_ = x, *ϕ*_3_ = y, *ϕ*_5_ = x, *ϕ*_7_ = y, *ϕ*_P_ = x and *ϕ*_S_ = y. In both SQ and ZQ/DQ experiments, the phase cycles are *ϕ*_2_ = (x, x, -x, -x), *ϕ*_4_ = (x, -x), *ϕ*_6_ = (y, -y) and *ϕ*_R_ = (x, -x, -x, x). For both experiments, the echo/anti-echo components are collected by incrementing *ϕ*_2_ and *ϕ*_6_ by 180° and by multiplying the encoding gradient amplitude by -1. All ^1^H and ^15^N hard pulses are broadband G5 composite pulses,^18^ and all gradients are 1.0 ms in duration. The Δ_1_, Δ_2_ and Δ_3_ periods are set to J_NH_^-1^/4 (2.78ms). The *δ*_1_ delay encompasses the durations of *δ*_1_’, t_1_ and the 180-degree ^13^C pulse length. The *δ*_2_’ delay encompasses the durations of *δ*_2_, t_1_, the 180-degree ^13^C pulse length, the 180-degree ^1^H pulse length and, if applicable, the *δ*_1_, *δ*_1_’ delays and an additional 180-degree ^1^H pulse length (ZQ/DQ). The encoding gradient field strength is 10.1% of the decoding gradient field strength.

In the ZQ/DQ experiment, MQ coherences remain after the initial insensitive nuclear enhancement through polarization transfer (INEPT) period by applying a ^1^H 90-degree pulse with an x-phase (*ϕ*_1_) instead of the y-phase used to generate the SQ coherence. Alternatively, this pulse could be removed in the ZQ/DQ experiment. Thereafter, the ZQ and DQ coherences are resolved by removing the blue 180-degree ^1^H pulse, which interchanges and averages the ZQ and DQ coherences in an HMQC.

The *δ*_1_/ *δ*_1_’ Hahn echo period ensures a 0°-per-point first order phase in the indirect dimension of the ZQ/DQ experiment. The *δ*_2_’/ *δ*_2_ Hahn echo period encapsulates the SQ encoding gradient, and it ensures a 0°-per-point first order phase in the indirect dimension of the SQ experiment.

### Spin physics

The free-induction decay (FID) intensities encode the chemical shifts and relaxation decays for the ^15^N-SQ, ZQ and DQ coherences.

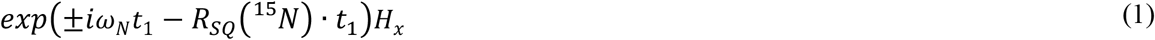

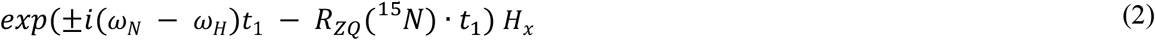

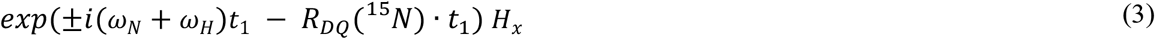

In the indirect (t_1_) dimension, the three types of coherences have relaxation rates that depend on ^1^H-^15^N dipolar and ^15^N chemical shift anisotropy (CSA) contributions.^10,11^

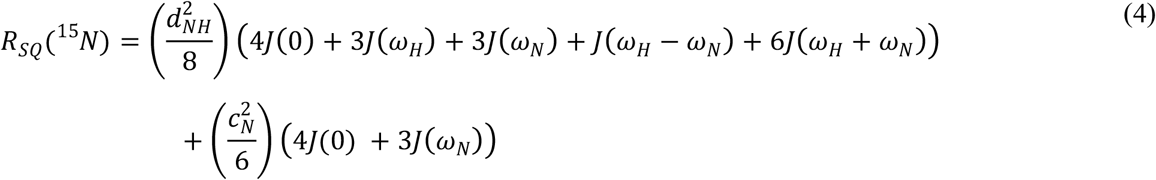

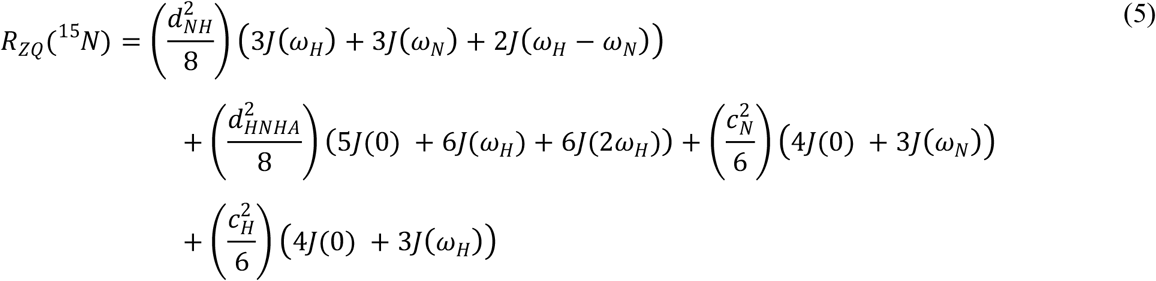

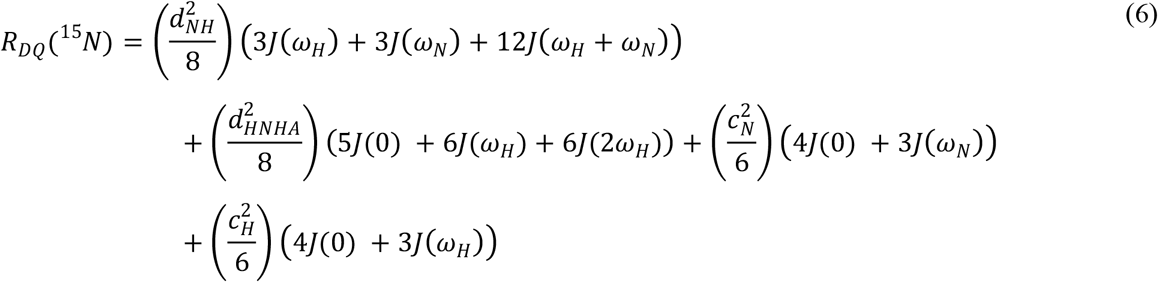

The R_SQ_(^15^N) relaxation rate is an average of the anti-phase 2H_z_N_+/-_ and in-phase N_+/-_ relaxation rates. The dipolar constant, d_NH_^2^ = (*µ*_0_/4*λ*)^2^ℏ^2^*γ*_H_^2^ *γ*_N_^2^r_NH_^-6^, and CSA constants, c_N_^2^ = *Δσ*_N_^2^*ω*_N_^2^/3 and c_H_^2^ = *Δσ*_H_^2^*ω*_N_^2^/3, are calculated from known constants. These constants depend on the vacuum permeability, *µ*_0_ = 1.2566 · 10^−6^ N/A^2^, the reduced Planck constant, ℏ = 1.055 · 10^−34^ Js, the ^1^H gyromagnetic ratio, *γ*_H_ = 267.522 · 10^6^ rad s^-1^ T^-1^, the ^15^N gyromagnetic ratio, *γ*_N_ = -27.116 · 10^6^ rad s^-1^ T^-1^, the H-N bond length, r_NH_ = 1.02 Å, the average ^15^N CSA for protein amides, *Δσ*_N_ = -160 ppm, and the average ^1^H CSA for protein amides, *Δσ*_H_ = 9ppm.^12,13^ The CSA constants and spectral densities are evaluated at the Larmor frequencies for ^1^H (*ω*_H_ = 2*λ*· 500MHz) and ^15^N (*ω*_N_ = 2*λ*· 50.7MHz) for a 500-MHz NMR spectrometer.

The d_HNHA_^2^ parameter represents the relaxation due to the dipolar coupling between the amide H^N^ and H_*α*_ protons. It is present for protonated proteins when the H^N^ magnetization is transverse in the indirect dimension of the ZQ and DQ transitions and in the direct dimension for all experiments. The d_HNHA_^2^ constant is fit as an average molecular parameter (discussed below), even though its value is expected to vary slightly as a function of the ϕ torsion angle. The SQ, ZQ and DQ transitions may also have contributions from chemical exchange, as discussed previously,^14^ and these are not modeled in this study.

The spectral densities, J(ω), can be fit using a variety of models.^15–17^ To minimize the simulation times in our spectral fits, we selected the simplest 2-parameter relaxation model.

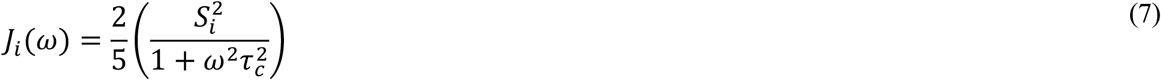

In this model, the spectral densities depend on the tumbling time for the molecule (*τ*_*c*_) and the generalized order parameter (Si^2^) for each resonance group (^1^H-^15^N pair).

The relaxation rate that governs the decay of H_x_ magnetization in the FID (direct dimension) is calculated as follows:

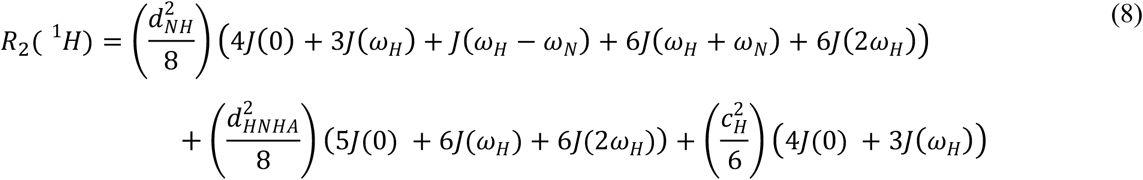

In the current implementation, the spectrum was collected with an indirect dwell time that encompasses SQ, ZQ and DQ frequencies (^15^N sweep width of *ca*. 80-100 ppm or a dwell time of *ca*. 200 *µ*s at 500-MHz). The shorter dwell time effectively doubles or triples the experiment length, in comparison to an HSQC with a ^15^N sweep width of 30 ppm. However, the shorter dwell time is equivalent to signal averaging. We additionally used G5-composite pulses for all ^1^H and ^15^N 180° pulses in order to excite resonances over a larger bandwidth.^18^

The final dataset is de-interleaved and processed as 2Ds, then these are extracted into HSQC, HZQC and HDQC spectra for combined fitting (See Figure S1).

## Results

### Simulations and data fitting

We developed a spectrum fitting software to identify resonance groups common to all spectra. The overall procedure was to simultaneously fit the intensities, chemical shifts and dynamic parameters for resonance groups (^1^H-^15^N pairs) in the HSQC, HZQC and HDQC spectra. The free parameters were minimized, and new resonance groups were discarded if there was not a statistical improvement in the χ^2^ fit of all 3 spectra, according to an F-test with a 95% confidence interval.

The free parameters were divided into 3 categories: molecular parameters, resonance group parameters and peak parameters. The molecular parameters were common to all resonance groups and peak types, and these include the tumbling time for the molecule (*τ*_c_), the average d_HHA_^2^ dipolar constant, a small MQ offset to increase DQ or decrease ZQ peak positions in the ^15^N dimension. The final minimized MQ offset typically ranged between 0-0.3 Hz (0.0-0.007 ppm). The resonance group parameters were common to all peak types associated to a resonance group. Each resonance group had its own generalized order parameter (S^2^) and, if it included peak types with J-couplings, the ^3^J_HNHA_-coupling parameter. The peak parameters only included the intensity of the peak, and these were fit independently for all 3 spectra. In a future implementation, we intend to further constrain the intensities between spectra.

Each resonance group had a peak type associated with each spectrum. Altogether, peak types for SQ, ZQ and DQ relaxation, as described in equations (1)-(8), were used to model the HSQC, HZQC and HDQC spectra, respectively. Each peak type included a ^3^J_HNHA_-doublet in the ^1^H dimension and, for the ZQ and DQ peak types, included a ^3^J_HNHA_-doublet in the ^15^N dimension. We modeled the peak in each dimension as a simple FID oscillation centered at the peak’s chemical shift frequency, an exponential decay corresponding to the dimension’s πR_2_ and, if applicable, a cosine-modulation pattern to model J-coupled doublets. Thereafter, each FID was apodized, zero-filled, and Fourier transformed to match the experimental spectrum’s processing parameters. The 1-dimensional spectra were then extended to 2-dimensions with a Kroenecker product. The simulated spectra were combined for all resonance groups, and the combined χ^2^ was evaluated against the experimental HSQC, HZQC and HDQC spectra.

The final refinement included an initial refinement step and a subsequent multi-step refinement to identify new resonance groups. In the initial refinement, we identified resonance groups from peak maxima in the contour plot. These initial peaks were used to detect the sub-spectral regions to speed up the simulation, and these resonances groups were then discarded after the first multi-step refinement. The multi-step refinement identified new resonance groups from 50 Monte-Carlo trials for the resonance group parameters, followed by a parameter minimization. The parameter minimization in each refinement step minimized the molecular parameters, selected the peak model types for each resonance group and minimized the parameters for the new resonance group, then the peak and molecular parameters for all resonance groups. This procedure was repeated 10 times. We found that increasing the number of multi-step refinements did not significantly improve the final *χ*^2^_min_.

Once the resonance groups were identified from the low-field (500-MHz) HCQC spectra, a high-field (500→900-MHz) HSQC spectrum was simulated with spectral widths, chemical shift frequencies and spectral densities at 900-MHz. The fit ^3^J_HNHA_-couplings were set to 0 Hz to produce singlet peaks for each resonance group.

### Spectral Comparisons

We selected two systems for the spectral comparisons: *α*-synuclein and the C-terminal domain of the influenza M1 capsid protein (M1C). *α*-synuclein was selected because its’ chemicals shifts and purification protocol are well characterized,^19,20^ and it includes 131 assignable peaks in its ^15^N-HSQC. We collected the *α*-synuclein spectra at 280K to increase the degree congestion in the spectra and to reduce hydrogen exchange rates.^21^ M1C is a large membrane protein system in DPC micelles, and it contains 132 assignable peaks in its ^15^N-HSQC.^22^ M1C was additionally selected because these samples suffer from degradation issues, which can be observed from the multiple C-terminal peaks, and we wanted to test a congested system with a variety of linewidths.

A total of 5 simulated 500→900-MHz HSQC spectra were simulated from the 500-MHz HCQC (see SI for refinement statistics), and the refinement with the lowest *χ*^2^_min_ is presented in the following plots. A comparison between an experimental 500-MHz HSQC, the simulated 500→900-MHz HSQC and an experimental 900-MHz HSQC are shown in **Figure 3**. Regions of interest with favorable enhancements for the 500→900-MHz HSQC are shown in **Figure 4** (Figure S1 includes assignments).

**Figure 3.**
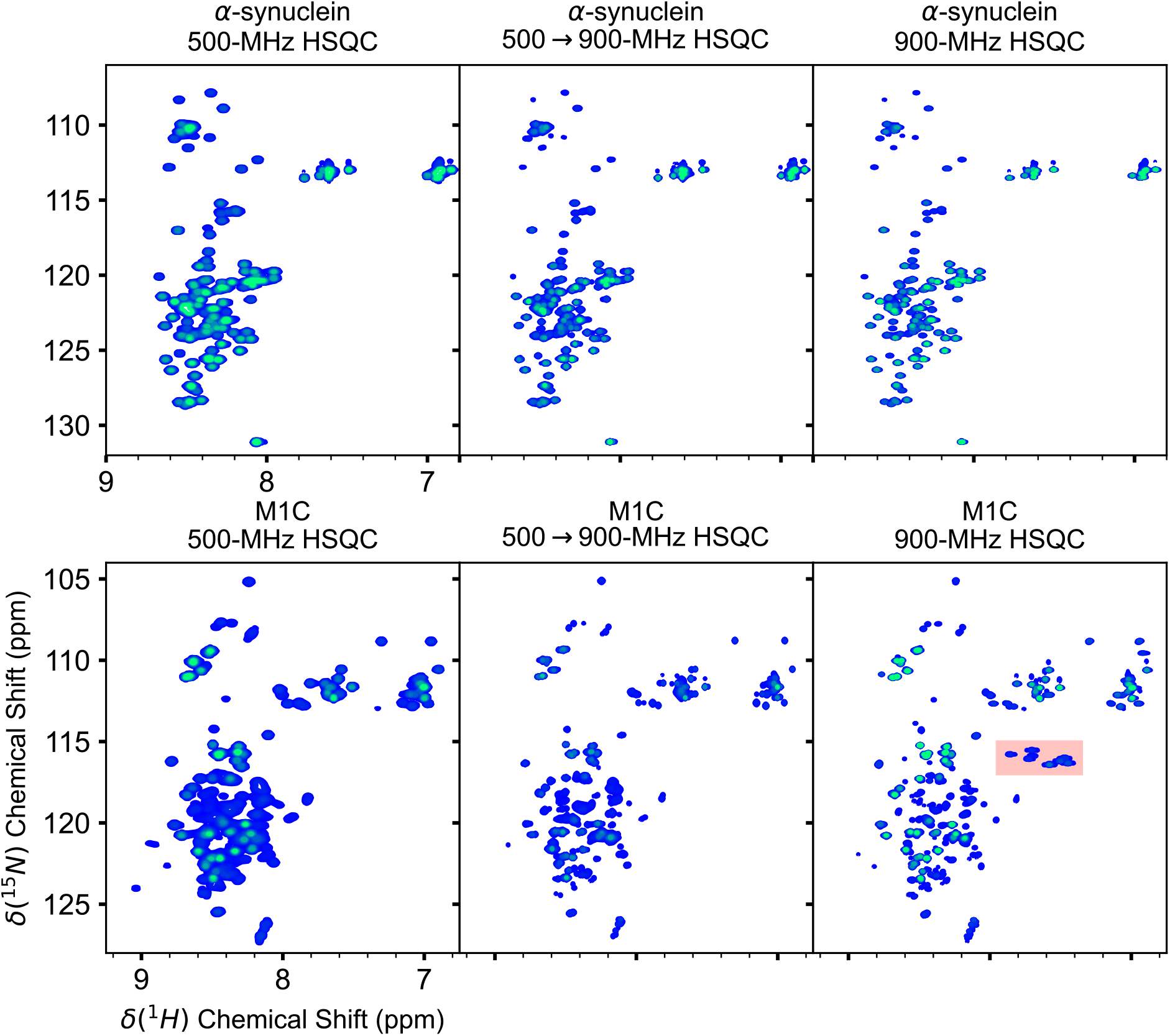
^15^N-HSQC 2D spectra for *α*-synuclein and M1C proteins. Spectra are shown for the experimental HSQC at 500-MHz, the simulated 500→900-MHz HSQC, and the experimental HSQC at 900-MHz. Contour levels for the *α*-synuclein spectra start at 4% of the intensity for the C-terminus, and contour levels for the M1C spectra start at 30% of the intensity for the ^15^N upfield glycine. The region marked in red in the M1C 900-MHz HSQC represent folded Arg sidechain resonances.

**Figure 4.**
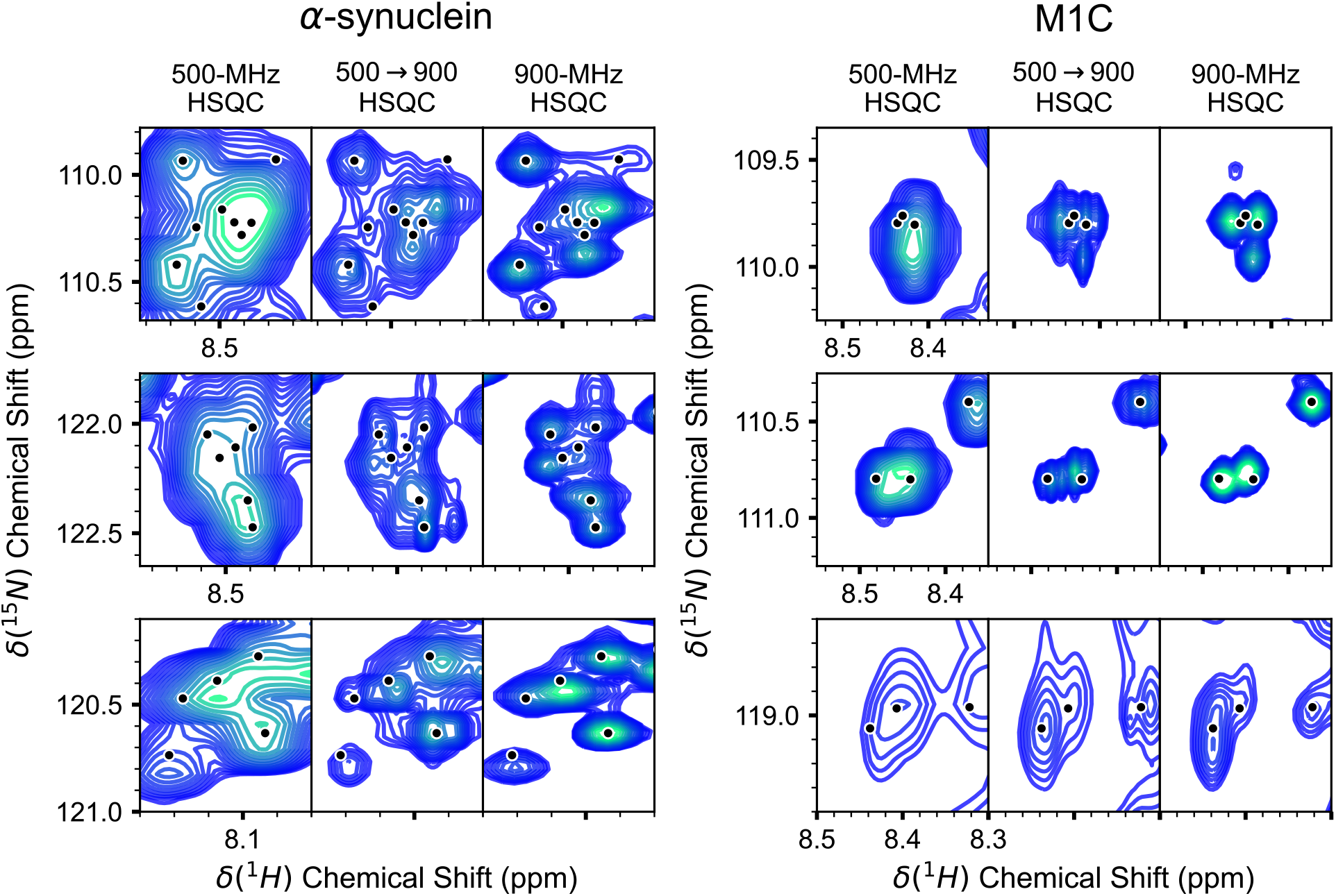
Selected regions of interest in the experimental 500-MHz HSQC, simulated 500→900-MHz HSQC and experimental 900-MHz HSQC spectra for α-synuclein (left) and M1C (right). Minor tick labels are separated by 0.02 ppm in the δ(^1^H) chemical shift dimension. See Figure S1 for assignment labels of peaks.

**Figure 5.**
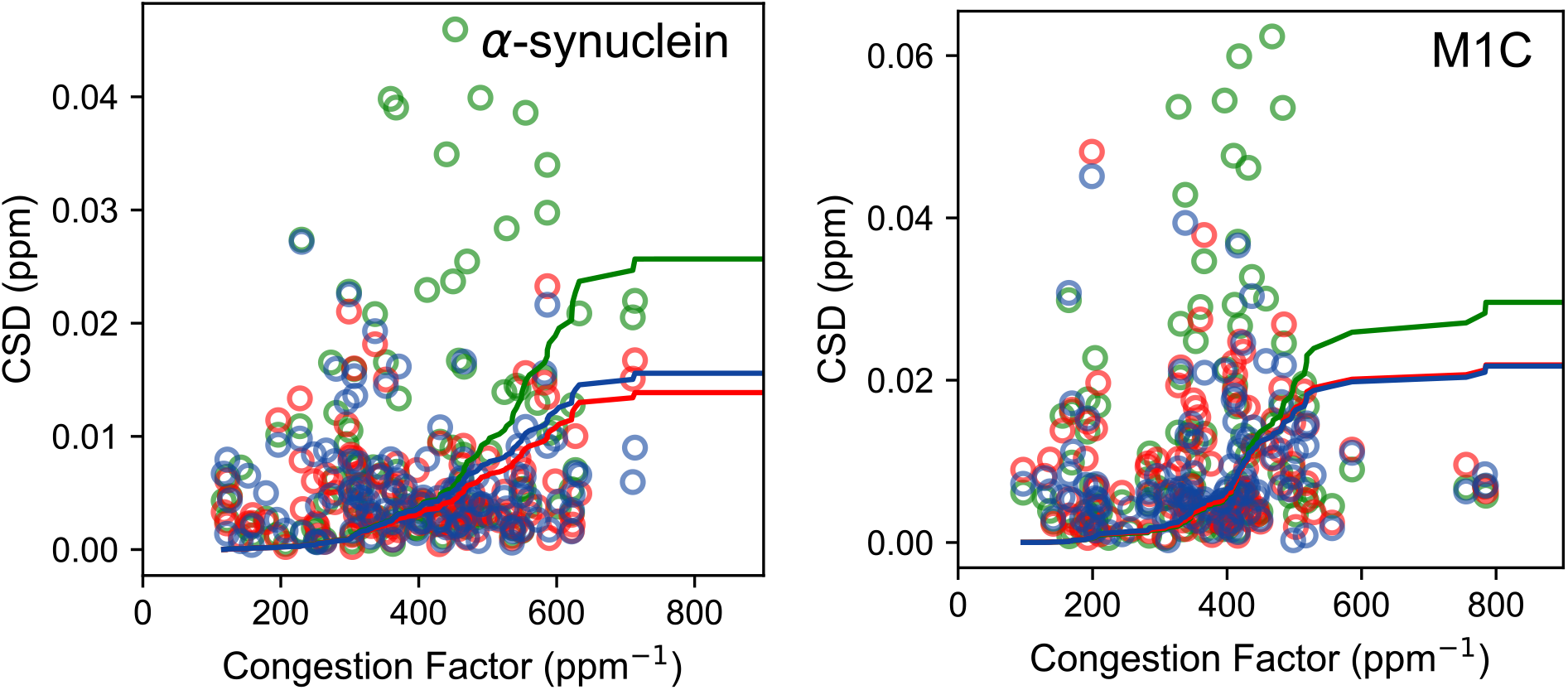
The chemical shift deviations (CSD) of assigned peaks in the HSQCs in comparison to the congestion factor. The chemical shift deviations are calculated from the ^1^H/^15^N peak positions of the 500-MHz HSQC (green points), the 500→900-MHz HSQC (blue points) and the 900-MHz HSQC (red points) in comparison to the closest assigned ^1^H/^15^N chemical shifts from a 500-MHz HNCO 3D. The lines represent the integration of the CSD, and these have been scaled by 40^-1^ in the plot. The integrated CSDs are 1.11, 0.67 and 0.60 ppm for the α-synuclein 500-MHz HSQC, 500→900-MHz HSQC and 900-MHz HSQC, respectively. The integrated CSDs are 1.42, 0.99 and 1.03 ppm for the M1C 500-MHz HSQC, 500→900-MHz HSQC and 900-MHz HSQC, respectively.

The *α*-synuclein peak positions and linewidths of the 500 →900-MHz HSQC closely match the experimental 900-MHz spectrum. The top row in the *α*-synuclein regions of interest show a portion of the glycine residues. The experimental 500-MHz HSQC shows 3 peaks (**Figure 4**, α-synuclein top row) whereas the 500 →900-MHz HSQC has 6 distinct peaks and the experimental 900-MHz has 6 distinct peaks. The central region has 4 peaks for residues G14, G47, G106 and G132, which merge into 1 peak in the 500-MHz HSQC. In the 500 →900-MHz HSQC, 4 peaks are observed whereas the 900-MHz HSQC only has 2-3 peaks in this central region. The G25 peak (8.52 and 110.6 ppm) can be seen in the 900-MHz and in the 500→900-MHz HSQC at lower contour levels. The G7 peak (8.45 and 109.9 ppm) is present in the 900-MHz HSQC but not the 500 →900-MHz HSQC. This difference could be attributable, at least in part, to the cryoprobe used to collect the 900-MHz spectrum in contrast to the room-temperature probe used to collect the 500-MHz spectra.

In the region containing residues Glu and Gln residues (**Figure 4**, α-synuclein middle row), *ca*. 1-2 peaks in the 500-MHz HSQC are resolved to *ca*. 6 peaks in the 500 →900-MHz HSQC and 7 peaks in the experimental 900-MHz. The resolved peaks in the 500→900-MHz HSQC and 900-MHz HSQC include E13, Q62, E105, E114 and E131, and the 900-MHz HSQC additionally resolves E61. The 500→900-MHz HSQC, however, includes a peak at 8.45 and 122.5 ppm in the lower right region, which is not observed in a 500-MHz HNCO 3D or the 900-MHz HSQC.

In the spectral regions containing Tyr and Phe residues (**Figure 4**, α-synuclein bottom row), 5 residues appear as 3-4 peaks in the 500-MHz HSQC. In the 500→900-MHz HSQC, these are resolved to 5 peaks for residues K32, F94, Y133, Y125 and Y136. In the 900-MHz HSQC, these are resolved to 4 peaks in which F94 and Y133 merge into the same peak.

The M1C system presents a much more challenging problem in correctly identifying the correct resonance groups in the spectrum. Multiple peaks are observed for C-terminal residues at *ca*. 8.1 ppm and 126.4 ppm (**Figure 3**). Most peak positions and linewidths are similar between the simulated 500→900-MHz HSQC and experimental 900-MHz spectra. Some differences are observed in regions that are resolved even in the experimental 500-MHz spectrum, such as the region at 8.8 ppm and 120 ppm.

These peaks and other peaks are present in the 500-MHz and 500→900-MHz HSQCs, but they are not visible at the contour levels of the plot, indicating that there are different relative intensities between spectra collected at 500- and 900-MHz.

Regions of interest in the M1C spectra show additional peaks resolved in the simulated 500→900-MHz HSQC spectrum. The glycine region (**Figure 4**, M1C top row) has 1 peak in the central region of the 500-MHz HSQC whereas theses are resolved to 3 peaks, G18, G31 and G96, in the 500→900-MHz HSQC and 900-MHz HSQC.

The glycine region encompassing residues G33 and G35 (**Figure 4**, M1C middle row) has 1 peak in the experimental 500-MHz HSQC. These 2 residues are resolved in the 500→900-MHz HSQC and 900-MHz HSQC. In this case, the simulated lineshapes are not accurately modeled in the spectrum: the central 2 peaks are fit to a singlet-doublet pattern with the base of each peak modeled with additional resonance groups.

The M1C region with residues N53, K106 and M124 (**Figure 4**, M1C bottom row) shows only 2 peaks in the 500-MHz spectrum. All 3 residues are resolved in the 500→900-MHz HSQC and 900-MHz HSQC. The 500→900-MHz HSQC additionally resolves a small contaminant peak for N53 at 8.32 and 119.0 ppm. This peak appears as a shoulder to the dominant N53 peak in the 500-MHz HNCO.

The black markers in **Figure 4** show the ^1^H and ^15^N chemical shift positions of peaks from a 500-MHz HNCO 3D experiment. Congestion even in a 3D spectrum presents challenges in accurately identifying the positions of peaks. For example, there are markers in the top and bottom row of the α-synuclein comparisons that are offset from the peak maxima in the 900-MHz HSQC and 500→900-MHz HSQC. This observation shows that the 500→900-MHz HSQC identifies peak positions more closely to the 900-MHz HSQC than the 500-MHz HNCO 3D spectrum.

We investigated the accuracy in the identification of new peaks in the 500→900-MHz HSQC using two methods—a manual peak assignment from known chemical shifts and an automated assignment based on assigned peak positions from the 500-MHz HNCO 3D spectrum. These were compared to conventional 500- and 900-MHz HSQC spectra.

### Manual peak assignment

The manual assignment of peaks is presented in Table 1 for *α*-synuclein and Table 2 for M1C. The total number of peaks (category 1) represents the number of singlets or J-coupled doublets from a simple peak picking procedure, using Sparky to identify peak maxima.^23,24^ Some doublets are observed in the 500→900-MHz HSQC, even though the ^3^J_HNHA_-coupling was set to 0 Hz for this spectrum. These represent peaks that were fit to 2 separate resonance groups a ^3^J_HNHA_-coupling smaller than the linewidth. Additionally, some simulated peak shapes fit better to an intense singlet with lower-intensity resonance groups at the base, forming a singlet-doublet pattern—for example, see the top and middle rows of the M1C 500→900-MHz HSQC spectrum in **Figure 4**. These simulated peak lineshapes are more complex than a simple singlet, but we counted these singlet-doublet patterns as individual peak groups.

**Table 1.**
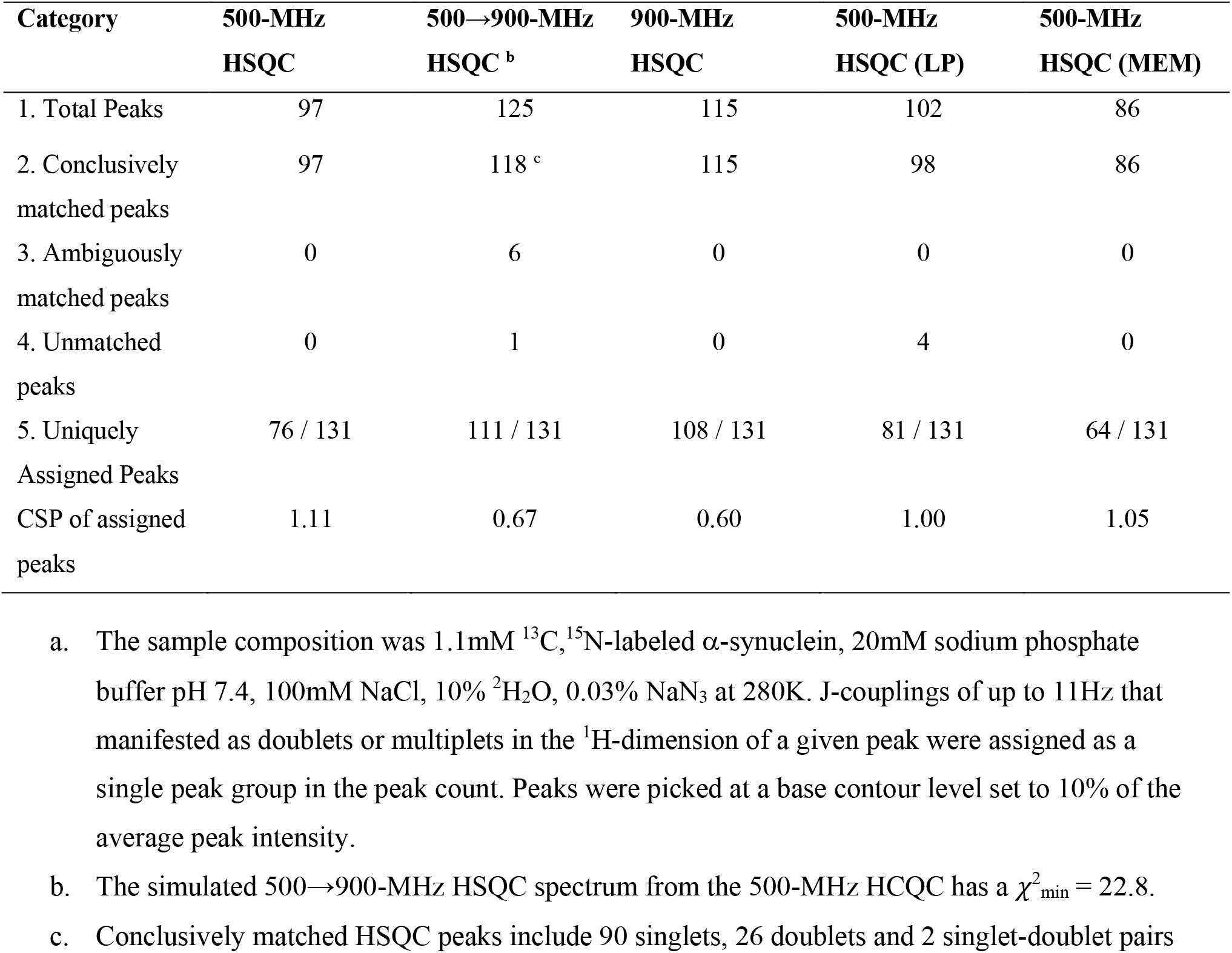
Spectral analysis of HSQC and HCQC spectral amide peaks for *α*-synuclein

| Category | 500-MHz<br>HSQC | 500→900-MHz<br>HSQC <sup>b</sup> | 900-MHz<br>HSQC | 500-MHz<br>HSQC (LP) | 500-MHz<br>HSQC (MEM) |
| --- | --- | --- | --- | --- | --- |
| 1. Total Peaks | 97 | 125 | 115 | 102 | 86 |
| 2. Conclusively<br>matched peaks | 97 | 118 <sup>c</sup> | 115 | 98 | 86 |
| 3. Ambiguously<br>matched peaks | 0 | 6 | 0 | 0 | 0 |
| 4. Unmatched<br>peaks | 0 | 1 | 0 | 4 | 0 |
| 5. Uniquely<br>Assigned Peaks | 76 / 131 | 111 / 131 | 108 / 131 | 81 / 131 | 64 / 131 |
| CSP of assigned<br>peaks | 1.11 | 0.67 | 0.60 | 1.00 | 1.05 |

a. The sample composition was 1.1mM <sup>13</sup>C, <sup>15</sup>N-labeled α-synuclein, 20mM sodium phosphate buffer pH 7.4, 100mM NaCl, 10% <sup>2</sup>H<sub>2</sub>O, 0.03% NaN<sub>3</sub> at 280K. J-couplings of up to 11Hz that manifested as doublets or multiplets in the <sup>1</sup>H-dimension of a given peak were assigned as a single peak group in the peak count. Peaks were picked at a base contour level set to 10% of the average peak intensity.
b. The simulated 500→900-MHz HSQC spectrum from the 500-MHz HCQC has a $\chi^2_{\min} = 22.8$ .
c. Conclusively matched HSQC peaks include 90 singlets, 26 doublets and 2 singlet-doublet pairs

**Table 2.** Spectral analysis of HSQC and HCQC spectral amide peaks for M1C ^a^

| Category | 500-MHz<br>HSQC | 500→900-MHz<br>HSQC <sup>b</sup> | 900-MHz<br>HSQC | 500-MHz<br>HSQC (LP) | 500-MHz<br>HSQC (MEM) |
| --- | --- | --- | --- | --- | --- |
| 1. Total Peaks | 92 | 144 <sup>c</sup> | 164 | 102 | 83 |
| 2. Conclusively<br>matched peaks | 88 | 138 | 156 | 95 | 80 |
| 3. Ambiguously<br>matched peaks | 4 | 6 | 8 | 7 | 3 |
| 4. Unmatched<br>peaks | 0 | 0 | 2 | 0 | 0 |
| 5. Uniquely<br>Assigned Peaks | 71 / 132 | 97 / 132 | 93 / 132 | 72 / 132 | 62 / 132 |
| CSP of assigned<br>peaks | 1.41 | 1.02 | 1.02 | 1.51 | 2.08 |

a. The sample composition was 1.2mM <sup>13</sup>C, <sup>15</sup>N-labeled M1C in 20mM sodium acetate, 150mM DPC, 10% <sup>2</sup>H<sub>2</sub>O, 0.03% NaN<sub>3</sub>, pH 5.01. All the spectra were collected at 305K. Peaks that show up as doublets or singlet-doublet pairs in HCQC spectrum due to J-coupling (<11 Hz) are counted as one peak group in the total peak count. Peaks were picked at a base contour level set to 10% of the average peak intensity.
b. The simulated 500→900-MHz HSQC spectrum from the 500-MHz HCQC has a $\chi^2_{\min} = 13.8$ .
c. Conclusively matched HSQC peaks include 121 singlets, 20 doublets and 3 singlet-doublet pairs

From the total number of peaks, peaks that can be matched to isolated peaks in a 500-MHz HNCO are identified as conclusively matched peaks (category 2). Peaks that matched congested regions in the HNCO are marked as ambiguously matched (category 3). Peaks in the HSQCs that could not be found in the 500-MHz HNCO 3D spectrum are marked as unmatched (category 4). Altogether, the number of peaks that can be uniquely assigned to one residue—as opposed to multiple residues—are counted as uniquely assigned (category 5).

For *α*-synuclein, the simulated 500→900-MHz HSQC has the largest number of peaks, 125, followed by the experimental 900-MHz HSQC, 115 peaks, and the experimental 500-MHz HSQC, 97 peaks. The simulated 500→900-MHz HSQC also has the most conclusively matched peaks, in comparison to the HNCO 3D spectrum, with 118 peaks. Overall, we achieve the largest number of unique assignments for the 500→900-MHz HSQC (111 of 131 possible assignments) in comparison to the experimental 900-MHz HSQC (108 of 131 assignments) and 500-MHz HSQC (76 of 131 assignments). The 500→900-MHz HSQC resolves the following 7 residues not resolved in the 900-MHz HSQC: K6, S9, G14, V49, G86 and F94. Conversely, the 900-MHz spectrum resolves 4 distinct peaks that appear as peak shoulders in the 500→900-MHz HSQC: G25, A27, Q79 and S87.

In comparison to existing resolution-enhancing strategies, linear prediction (LP) and maximum entropy reconstruction (MER), the 500→900-MHz HSQC spectrum performs significantly better. Altogether, LP processing only yielded 5 additional assignment and MER processing yielded 12 fewer assignments in comparison to the experimental 500-MHz HSQC. These procedures are limited in their resolution enhancements since they narrow linewidths or model peak shapes from the existing information in the 500-MHz HSQC. By contrast, the 500-MHz HCQC includes additional non-redundant information, which is used to more accurately model peak positions and to quantitatively increase the spectrum’s resolution.

The manual peak identification and assignment for M1C show a similar trend (Table 2) in the total number of peak assignments. The 900-MHz HSQC has the largest number of total peaks in comparison to the 500-MHz HSQC and simulated 500→900-MHz HSQC. The 900-MHz HSQC was collected on a cryoprobe whereas the 500-MHz HSQC and 500-MHz HCQC were collected on a room-temperature 500-MHz NMR spectrometer. Consequently, additional minor peaks for contaminants are observed in the total number of peaks and the number of conclusively matched peaks in the 900-MHz HSQC.

We nevertheless can assign the greatest number of peaks in the 500→900-MHz HSQC (97 of 132 possible assignments) in comparison to the experimental 500-MHz HSQC (71 of 132 assignments) and 900-MHz HSQC (93 of 132 assignments). We identify 5 residues in the 500→900-MHz HSQC— residues S11, S12, G22, G39 and M124—not resolved in the 900-MHz HSQC. Conversely, we identify 1 additional residue in the 900-MHz HSQC—G96—but not the 500→900-MHz HSQC.

### Automated peak assignment

We conducted an automated peak assignment strategy as an alternative to manual assignment. For this strategy, we used the default peak-picking algorithm in Sparky^23,24^ to identify peaks in the 500-MHz HSQC, 500→900-MHz HSQC, 900-MHz HSQC and the 500-MHz HNCO 3D. Thereafter, the peaks from the reference HNCO 3D were matched to the closest peak found in each HSQC and the chemical shift difference (CSD) between peak positions were calculated as follows:

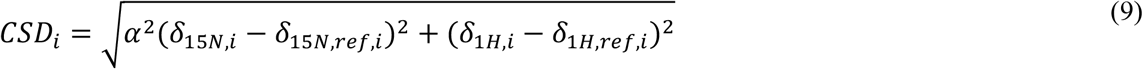

In this study, we used an α value of 0.10.

We expected the CSDs to be excellent for isolated peaks and to be poor for congested peaks. Consequently, we plotted the CSDs as a function of a chemical shift congestion factor (CF). The CF measures the sum of ^1^H/^15^N pairwise distances of a peak in the HNCO to all other peaks.

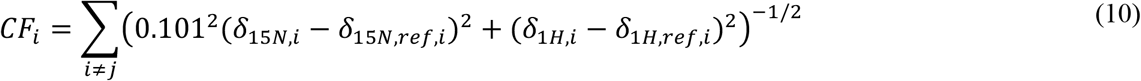

Peaks with small congestion factors are present as isolated peaks in the HSQCs whereas peaks with large congestion factors are present in highly congested or overlapped regions in the HSQCs.

Since this analysis depends on the chemical shift referencing of the spectra, the overall CSD was minimized for the ^1^H/^15^N offset of each HSQC spectrum in comparison to the HNCO 3D spectrum’s chemical shifts. We found that the referencing correction was < 0.003 ppm in the ^1^H dimension and < 0.055 ppm in the ^15^N dimension for all HSQCs.

The inaccuracy of peak positions in the 500-MHz HSQC is most apparent in regions with a congestion factor of *ca*. 250-700 ppm^-1^ and above. The overall, integrated CSD for *α*-synuclein is 1.11 ppm, 0.67 ppm and 0.60 ppm for the 500-MHz HSQC, 900-MHz HSQC and 500→900-MHz HSQC, respectively. The accuracy of peak positions for the 500→900-MHz HSQC is not quite as high as the 900-MHz HSQC with a reduction of only 86% of the difference between the overall CSD in the 500-MHz and 900-MHz HSQCs. By contrast, the 500-MHz HSQC with LP and MER processing have overall CSDs of 1.00 ppm and 1.05 ppm, which is nearly identical to the reference 500-MHz HSQC (Table 1).

The M1C CSDs show a similar trend. The overall, integrated CSD is 1.42, 0.99 and 1.03 ppm for the 500-MHz HSQC, 500→900-MHz HSQC and 900-MHz HSQC respectively. The overall CSD for LP processing is 1.50 ppm and for MER processing it is 2.08 ppm, which are comparable or worse to the reference 500-MHz HSQC. Altogether, the peak positions of the 500→900-MHz HSQCs are nearly comparable in accuracy to the 900-MHz HSQCs.

## Discussion

INIR spectra offer significant resolution enhancements. The number of assignments and peak positions from a reconstructed 500→900-MHz HSQC spectrum approach the accuracy and coverage of a 900-MHz HSQC spectrum, which represents a spectrometer with an 80% stronger magnetic field strength. The current implementation nevertheless suffers from a few disadvantages.

The first limitation is in the accurate modeling of peak shapes in the current simulation protocol. The number of fit resonance groups exceeds the number of assignable resonances by a factor of 2-3 (Table S1 and S2). Each resonance group can be fit to (1) a doublet in the ^1^H dimension, separated by the ^3^J_HNHA_-coupling, (2) one or more singlets in which the ^3^J_HNHA_-coupling is smaller than the peak’s ^1^H linewidth, or (3) a high-intensity peak with reduced intensity resonance groups at the base of the peak. We included the impact of ^3^J_HNHA_ couplings in the simulation because many peaks manifest as clear doublets in the ^1^H dimension and these couplings impact the ZQ and DQ linewidths in the HZQC and HDQC, respectively. However, the variety of possible peak shapes can present multiple minima in the *χ*^2^ refinement. An alternative pulse sequence could remove these effects by using a selective ^1^H^*α*^ inversion pulse in the HZQC and HDQC and by applying ^1^H homonuclear decoupling in the direct dimension.^25,26^ Additionally, perdeuteration of the molecule would suppress these couplings in the spectrum. We elected not to suppress ^3^J_HNHA_-couplings in this first application to test the technique’s amenability for biomolecules with only ^15^N isotope labels and to keep the pulse sequence simple. Further improvements to the experiment and simulation strategy, such as the inclusion of Voigt lineshape profiles and the removal of redundant resonance groups, would nevertheless improve the accuracy and speed of our refinements.

The impact of the molecule’s tumbling time, *τ*_c_, and the resonance group order parameter, S^2^_i_, are other parameters which can impact the accuracy of peaks fits. The *τ*_c_ estimates from the best-fit refinement range between 5.0-7.1 ns for *α*-synuclein (Table S1) and 13.5-18.2 ns for M1C (Table S2). The experimentally determined *τ*_c_ for *α*-synuclein is 9.5 ± 2.5 ns at 288K ^27^ and our experimentally determined *τ*_c_ for M1C is 13.7 ± 0.1 ns at 305K from ^15^N relaxation studies.^22^ Shorter tumbling times, smaller order parameters or a combination of both will yield narrower peaks, and complex experimental peak lineshapes may be fit with lower *χ*^2^ values to a combination of multiple narrow peaks. Constraining these parameters could further simplify the modeling of peaks.

The second limitation is the additional experimental time and simulation time required for INIR spectra. At least 2 interleaved spectra are required and, in the case of the HZQC and HDQC spectra, a shorter dwell time is required in the indirect dimension. Our experimental HCQC 2D spectra took at least 8 hours to collect. In principle, the collection of HCQC spectra is complementary with NUS collection,^28,29^ and HCQC spectra could be collected in a much shorter time. The computational time to refine spectra is also quite long at *ca*. 24 hours per refinement. This shortcoming could be overcome by developing code that does not rely on an interpreted language (Python) for computationally-intensive operations and to use graphics processing units (GPUs) for the minimizations.

Despite the shortcomings, INIR spectroscopy offers important advantages in the analysis of NMR spectra. First, the required dimensionality, experimental acquisition time and complexity of spectra may be reduced. For large biomolecular systems, conventional experiments may require 3D or 4D experiments to ascertain experimental values for most or all spin systems. For example, KcsA is a membrane protein system with 160 residues, and 4D relaxation TROSY-HNCO experiments are required to identify the ^15^N-R_1_, ^15^N-R_2_ and {^1^H}-^15^N NOE relaxation rates. ^30^ Enhanced resolutions could enable simplified 3D relaxation experiments based on HSQCs. Spectra that require additional labeling strategies, such as uniform ^13^C labeling, may be circumvented as well, and the utility of expensive, high-field NMR spectrometers may be improved.

Second, combinations of other interleaved spectra could be refined together to model the parameters of resonance groups more accurately. Examples could include Hahn echos to resolve inhomogeneous broadening, diffusion-ordered spectroscopy (DOSY) to resolve different molecules and ^15^N-relaxation experiments that, together with the SQ, ZQ and DQ transitions, more accurately deconvolve clusters of resonance groups simultaneously. Additionally, the incorporation of R_ex_ terms in the SQ, ZQ and ZQ transitions would give additional information on the microsecond-to-millisecond dynamics of the system.^14^

Third, our application of INIR spectroscopy could be extended to higher dimensional experiments, such as HNCO, HNCA, HNCOCA experiments.^31^ An INIR HNCO 3D, which would require 2-4 times the experiment time to collect, could be used to further resolve spins in a 3D at low field. This experiment is advantageous at low field since carbonyl resonance linewidths are dominated by the CSA, which increases quadratically with field strength, whereas the chemical shift dispersion only scales linearly with field strength.

Fourth, enhanced resolution presents an opportunity to study larger biomolecular systems. The molecular size of systems amenable for solution NMR study is limited by coherence lifetimes and spectral congestion. Larger molecules have larger rotational tumbling times, *τ*_c_, thereby reducing coherence lifetimes and increasing peak widths. Bigger molecules also have more resonance groups that further congest the spectrum. Enhancing the resolution of spectra would enable the analysis of more complex and larger biomolecules.

## Conclusions

We have demonstrated an initial application of INIR to enhance the resolution of NMR spectra. The current implementation is useful as a guide to identify new peaks in a spectrum. Further developments hold promise in producing a quantitative, high-resolution reconstruction of a spectrum and in enabling a wider range of applications for low-field and high-field NMR spectrometers. These developments include improvements in the identification and simulation of resonance groups, the integration of alternative interleave experiments, and the exploration of isotope labeling schemes to improve resolution.

## Acknowledgements

J.L.L and I.B. were supported by NSF grant #MCB1651598. This study made use of the National Magnetic Resonance Facility at Madison, which is supported by NIH grant R24GM141526. We thank Prof. Ann E. McDermott for reviewing a draft of this manuscript and for her helpful comments.

## Materials and Methods

### α-synuclein sample preparation

The pET-41 cDNA plasmid vector that encoded human α-synuclein with a T7 lac promoter (generously provided by the Ad Bax group) and kanamycin resistance was used to transform *E. coli* BL21 (DE3) competent cells. The overnight cultures were grown in 50 mL of M9 minimal media supplemented with 1.5 g ^15^NH_4_Cl (Cambridge Isotope Laboratories) and ^13^C-D-glucose (Cambridge Isotope Laboratories) for ^13^C/^15^N-labeled protein at 37°C and 200rpm.^32^ The overnight cultures were used to inoculate the remaining 950mL of minimal media. The cells were grown to an optical density at 600nm (OD_600_) of ∼0.7 and induced with 1 mM IPTG (Gold-Bio, St. Louis, MO, USA) for a period of 5 hours before harvesting at 8,000x g and 4°C for 20 minutes. The supernatant was discarded, and the pellet was resuspended and washed in 30mL of milliQ H_2_O, centrifuged at 4,500x g and 4°C for 30 minutes, the supernatant was then discarded and the pellet was stored at -80°C until purification.

The cell pellet was resuspended in 10mL of Lysis Buffer [50mM Tris pH 7.4, 500mM NaCl, 0.5mM EDTA] on an orbital shaker until thawed. Resuspended cells were subjected to heat precipitation at 90°C in 5-minute intervals for 25 minutes with 30 seconds of vortexing between intervals. The mixture was diluted to 100 mL with Q-Sepharose Buffer A [20mM Tris pH 7.4, 0.5mM EDTA], the mixture was then centrifuged at 75,000x g and 4°C for 30 minutes. The supernatant was injected onto a Q-sepharose column equilibrated to 12% Buffer B [20mM Tris pH 7.4, 500mM NaCl, 0.5mM EDTA] then eluted with 24% Buffer B. Immediately upon elution, the fraction containing the protein was injected to a HiLoad 26/600 Superdex 200 pg (Cytiva) and eluted with the Run Buffer [20mM Tris pH 7.4, 150mM NaCl, 0.5mM EDTA].

Fractions of monomeric α-synuclein were combined and concentrated in a 20mL, 10kDa MWCO centrifugal concentrator (Vivaspin) until the total volume was less than 1mL. NMR Buffer [20mM NaPi Buffer pH 7.4, 100mM NaCl, 10% D_2_O, 0.03 NaN_3_] was serially aliquoted to the concentrator in 5mL increments and centrifuged at 4500x g and 4°C until the total volume was less than 1mL. The process was repeated four times, and on the fifth time the solution was concentrated to 1 mL, and the final concentration was measured with a NanoDrop 2000 (Thermo Scientific) using the extinction coefficient, 5120 M^-1^cm^-1^, and average molecular weight, 14460.16 Daltons, then split to two separate NMR tubes and measured within 48 hours of preparation. The final composition of the NMR samples consisted of 1.1mM ^13^C,^15^N-labeled α-synuclein in NMR Buffer [20mM sodium phosphate buffer pH 7.4, 100 mM NaCl, 10% ^2^H_2_O, 0.03% NaN_3_], and experiments were measured at 280K (7°C).

### M1C sample preparation

The C terminal domain of Influenza A Matrix protein 1 (M1C) was subcloned into pET15b vector (GenScript). In addition to the M1C sequence, the construct contained a hexa-histidine tag, a Strep-II tag and a Factor Xa cleavage site on the N-terminus and a GSKKKKD solubility tail on the C-terminus. The recombinant plasmid was transformed into *E. coli* BL21 (DE3) cells for protein expression and purification. M1C with uniform ^13^C and ^15^N isotope labels was expressed in 1L M9 minimal media containing 1.8 g ^15^NH_4_Cl and 2.0 g ^13^C-glucose (Cambridge Isotope Laboratories). Cells were grown at 37°C, 200 rpm and induced with 1 mM IPTG (GoldBio, St.Louis, MO, USA). Cells were harvested 4 hours after induction when the optical density (OD_600_) reached a value of 0.8. Cells were harvested by centrifucation at 8000x g for 30 min at 4°C, and the cell pellet was stored at -80°C until purification.

The cell pellet was resuspended in 50-100mL Lysis Buffer [8M urea (Spectrum Chemical, New Brunswick, NJ, USA), 50mM Tris, 500mM NaCl, 20mM Imidazole pH 8.0]. The cells were then sonicated to produce a clarified solution and centrifuged at 70,000x g for 30 min at 4°C. The supernatant was loaded to a 5mL HisTrap column (Cytiva, Marlborough, MA, USA) attached to an AKTA Start FPLC system (Cytiva). The column was washed with approximately 100 mL of Lysis Buffer and bound protein was eluted using a linear gradient of Elution Buffer [8M urea, 50mM Tris, 500mM NaCl, 500mM Imidazole pH 8.0]. The eluted protein was dialyzed in 4L water at 4°C overnight. The dialyzed protein was further purified using Reverse Phase Chromatography (RPC). We used a 3 ml Resource RPC column (Cytiva) on an AKTA Purifier (Cytiva) system. The dialyzed protein containing 5% Acetonitrile (Fisher Scientific, Hampton, NH, USA) and 0.1% TFA was injected to the pre-equilibrated column and eluted with a 10 min gradient to 80% acetonitrile/0.1% TFA. The purified protein was lyophilized before further use.

The M1C NMR samples were prepared by solubilizing lyophilized protein to a final concentration of 1.2 mM ^13^C,^15^N-labeled M1C in NMR Buffer [20mM Sodium Acetate, 150mM DPC (Anatrace, Maumme, OH, USA), 10% ^2^H_2_O, 0.03% NaN_3_ at pH 5.0], and experiments were collected at 305K (32°C).

### NMR experiments

Spectra were collected on a 500-MHz Bruker Avance III NMR spectrometer with a 5mm ^1^H/^2^H/^13^C/^15^N/^31^P QXI room-temperature probe and a 900-MHz Bruker Avance III HD NMR spectrometer at the University of Wisconsin Madison with a 5mm ^1^H/^2^H/^13^C/^15^N TXI cryoprobe. Reference HSQC 2D spectra were collected with the ‘hsqcetfpf3gpsi2’ pulse sequence,^3–5,33^ an ^15^N acquisition time of 120 ms, a ^1^H acquisition time of 80 ms and a recycle delay of 1.3-1.5 s. The reference HNCO 3D spectra were collected with the ‘hncogp3d’ pulse sequence, an ^15^N acquisition time of 22.4 ms, a ^13^C acquisition time of 19.5-23.1 ms, a ^1^H acquisition time of 80 ms and a recycle delay of 1.3-1.5 s.

The HCQC experiment was implemented as described in the main text. The pulse sequence was based on the ‘hsqcetf3gpsi’ pulse sequence and included an interleaved HSQC with sensitivity improvement and an HMQC with sensitivity improvement modified by removing the ^1^H 180° refocusing in the middle of the t_1_ period. The 180° pulses were implemented as composite G5 pulses to refocus or invert magnetization over a larger frequency range.^18^

### Spectral simulations

A simulation software ‘specfit’ was developed to simultaneously fit HSQC, HZQC and HDQC spectra from HCQC experiments. The software was developed using Python 3.9 and the following major libraries: numpy^34^ and scipy^35^ for numerical operations, lmfit for minimization,^36^ opt_einsum^37^ for Kroenecker products, and nmrglue for data conversion. ^38^ Details on the simulation protocol are described in the Results. The source code for specfit can be found at https://github.com/jlorieau/specfit.

Spectra and plots were prepared with matplotlib.^39^

## Supplementary Information

**Figure S1.**
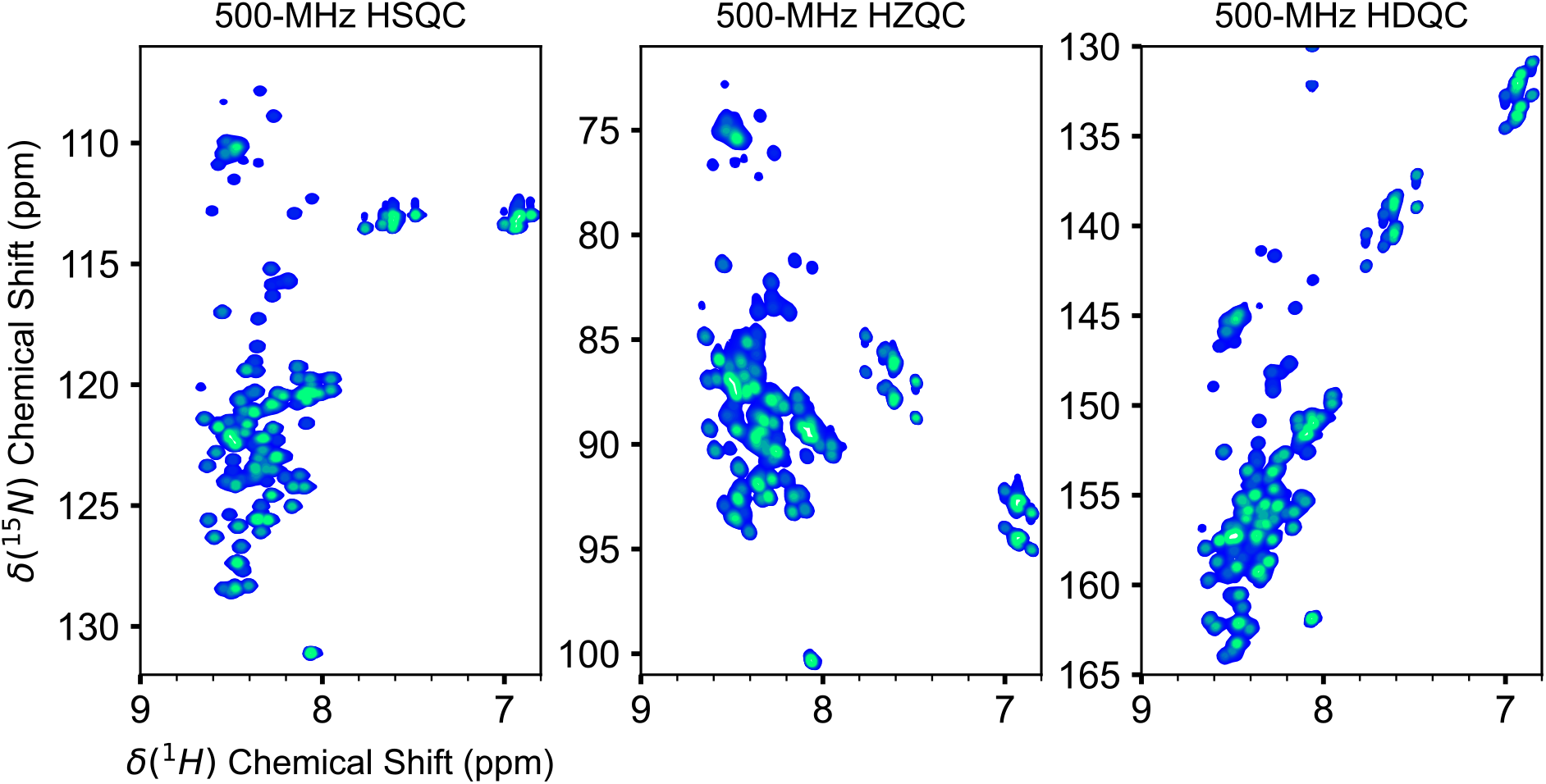
The separated HSQC, HZQC and HDQC from a 500-MHz HCQC of synuclein collected at 280K.

**Figure S2.**
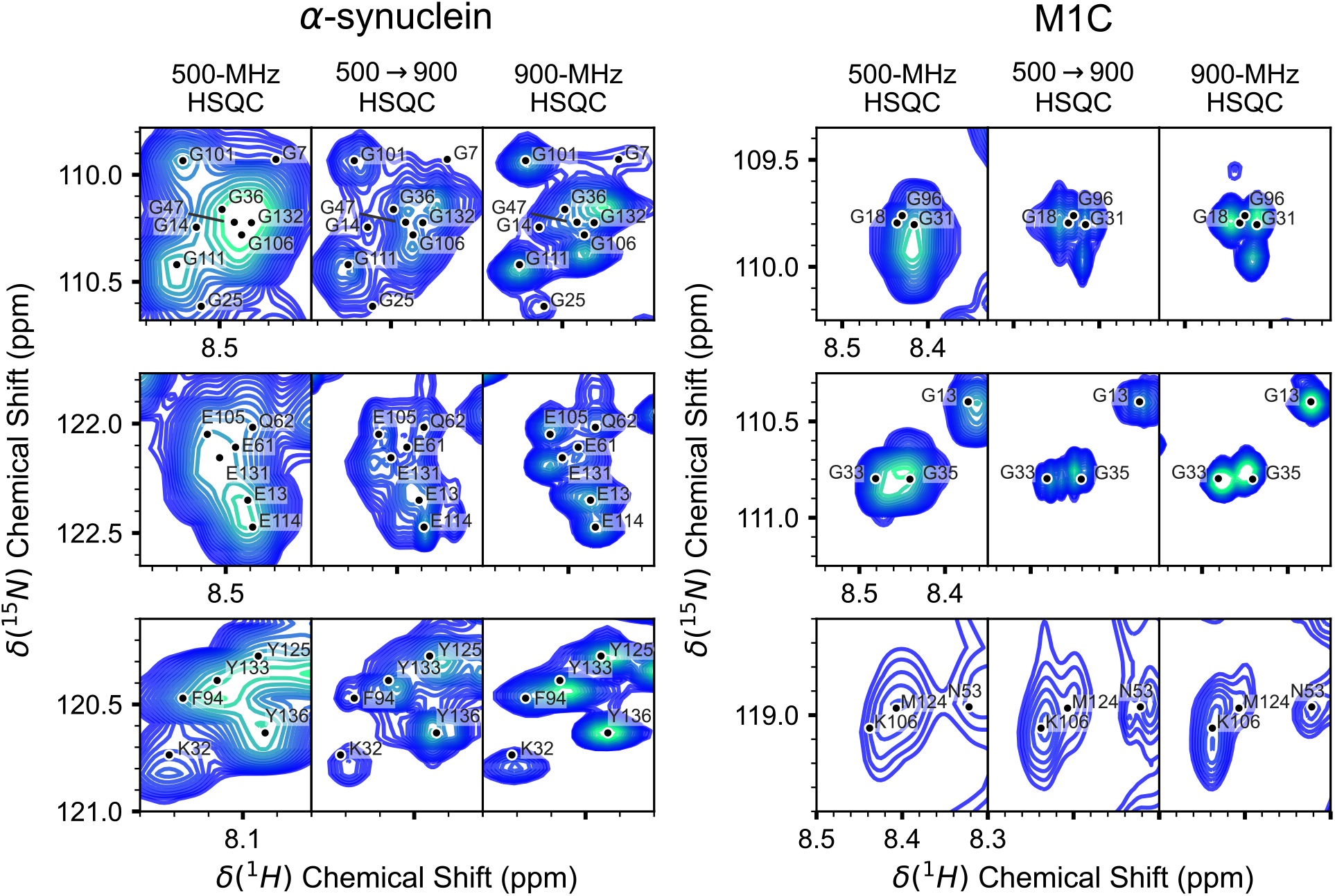
Selected regions of interest in the 500-MHz, 500→900-MHz simulated HSQC and 900-MHz HSQC spectra for α-synuclein (left) and M1C (right), Figure 4, with assignment labels.

**Table S1.** Simulated 900-MHz HSQC spectra datasets for *α*-synuclein reconstructed from a 500-MHz HCQC

| Run | $\chi^2_{\min}$ | Number of Resonance<br>Groups<sup>a</sup> | Estimated $\tau_c$ (ns) | CSP (ppm) |
| --- | --- | --- | --- | --- |
| 1 | 22.8 | 339 | 6.1 | 0.665 |
| 2 | 23.1 | 328 | 7.6 | 0.730 |
| 3 | 23.9 | 294 | 5.0 | 0.706 |
| 4 | 25.1 | 323 | 6.1 | 0.725 |
| 5 | 26.5 | 389 | 6.5 | 0.701 |

a. Peaks with a SNR > 10

**Table S2.** Simulated 900-MHz HSQC spectra datasets for influenza M1C reconstructed from a 500-MHz HCQC

| Run | $\chi^2_{\min}$ | Number of Resonance<br>Groups <sup>a</sup> | Estimated $\tau_c$ (ns) | CSP (ppm) |
| --- | --- | --- | --- | --- |
| 1 | 13.8 | 388 | 16.9 | 0.991 |
| 2 | 15.1 | 285 | 13.5 | 1.099 |
| 3 | 15.6 | 365 | 17.6 | 1.119 |
| 4 | 15.9 | 380 | 18.2 | 1.082 |
| 5 | 16.4 | 313 | 15.5 | 1.184 |

b. Peaks with a SNR > 10

